# SPAligner: Alignment of Long Diverged Molecular Sequences to Assembly Graphs

**DOI:** 10.1101/744755

**Authors:** Tatiana Dvorkina, Dmitry Antipov, Anton Korobeynikov, Sergey Nurk

## Abstract

**Background:** Graph-based representation of genome assemblies has been recently used in different applications — from gene finding to haplotype separation. While most of these applications are based on the alignment of molecular sequences to assembly graphs, existing software tools for finding such alignments have important limitations.

**Results:** We present a novel SPAligner tool for aligning long diverged molecular sequences to assembly graphs and demonstrate that SPAligner is an efficient solution for mapping third generation sequencing data and can also facilitate the identification of known genes in complex metagenomic datasets.

**Conclusions:** Our work will facilitate accelerating the development of graph-based approaches in solving sequence to genome assembly alignment problem. SPAligner is implemented as a part of SPAdes tools library and is available on https://github.com/ablab/spades/archive/spaligner-paper.zip.

## Background

Many popular short read assemblers [1, 2, 3] provide the user not only with a set of contig sequences, but also with assembly graphs, encoding the information on the potential adjacencies of the assembled sequences. Naturally arising problem of sequence-to-graph alignment has been a topic of many recent studies [4, 5, 6, 7, 8]. Identifying alignments of long error-prone reads (such as Pacbio and ONT reads) to assembly graphs is particularly important and has recently been applied to hybrid genome assembly [9, 10], read error correction [11], and haplotype separation [12]. At the same time, the choice of the practical aligners supporting long nucleotide sequences is currently limited to vg [4] and GraphAligner [13], both of which are under active development. Moreover, to the best of our knowledge, no existing graph-based aligner supports alignment of amino acid sequences.

Here we present the SPAligner (Saint Petersburg Aligner) tool for aligning long diverged molecular (both nucleotide and amino acid) sequences against assembly graphs produced by the popular short-read assemblers. The project stemmed from our previous efforts on the long-read alignment within the hybridSPAdes assembler [9]. Our benchmarks on various Pacbio and Oxford Nanopore datasets show that SPAligner is highly competitive to vg and GraphAligner in aligning long error-prone reads. We also demonstrate SPAligner’s ability to accurately align amino-acid sequences (with up to 90% amino acid identity) onto complex assembly graphs of metagenomic datasets. To further motivate this application we show how SPAligner can be used for identification of biologically important (antibiotic-resistance) genes, which remain under the radar of conventional pipelines due to assembly fragmentation (e.g. genes exhibiting high variability in complex environmental samples).

## Implementation

Section “Alignment of long nucleotide sequences introduces” describes the notation and presents the approach taken in SPAligner for semi-global alignment of nucleotide sequences to the assembly graph. Extension of this approach for aligning amino acid sequences is discussed in Section “Alignment of amino-acid sequences”.

### Alignment of long nucleotide sequences introduces

Let *G* be a (directed) compacted de Bruijn graph with edges labeled by nucleotide sequences that we further refer to as assembly graph ^1^. Length of a nucleotide sequence *S* is denoted by |*S* |and length of edge *e*, |*e*|, is defined as a length of its label. *S*[*x*] denotes *x*-th symbol of *S* (starting from 0). Position *x* on string *S* belongs to a range [0 … |*S*|] and corresponds to location before *S*[*x*], if *x* < |*S*|, and after *S*[|*S*| − 1] otherwise. *S*[*a*: *b*] is a substring of *S* between positions *a* and *b*. By the size of graph *G*, |*G*|, we denote the sum of the number of its vertices, edges and total length of all labels.

Position in the graph is naturally defined by a pair of an edge *e* and position in the sequence of *e*: *p* = (*e, i*), where 0 *<*= *i* < = |*e*|. Note that with such notation there are multiple positions in graph that corresponds to a vertex of *G* — it is located at the end of each incoming edge and at the beginning of each outgoing edge. *G*[*p*] denotes a symbol on position *i* within edge *e*. A path *P* in *G* is defined by a sequence of consecutive edges *e*_1_, …, *e*_*n*_ and a pair of positions 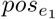 and 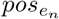 on *e*_1_ and *e*_*n*_ respectively. We denote the nucleotide sequence obtained by concatenation of edge labels in a path *P* (trimmed according to 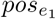 and 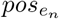) as *Label*(*P*).

For the sake of clarity throughout the paper we are going to neglect the fact that adjacent edges produced by popular assembly tools typically overlap (in particular adjacent edges produced by majority of the de Bruijn graph based assemblers overlap by *K* bp, where *K* is the kmer size parameter of de Bruijn graph).

The semi-global alignment of a sequence *S* (query) to an assembly graph *G* is defined as a path *P* in *G*, such that *Label*(*P*) has minimal alignment cost against *S* across all paths in *G*. As well as other practical tools for searching alignments in large sequence graphs (including vg and GraphAligner) SPAligner first identifies regions of high nucleotide identity between the query and the graph sequences that we refer to as anchor alignments (or anchors) and then attempts to extend them to desired semi-global alignments.

Below we outline the four steps implemented in SPAligner for aligning a nucleotide sequence query *S* to an assembly graph *G* (see Figure 1), generally following the approach used in alignment module of hybridSPAdes assembler [9]

**Figure 1:**
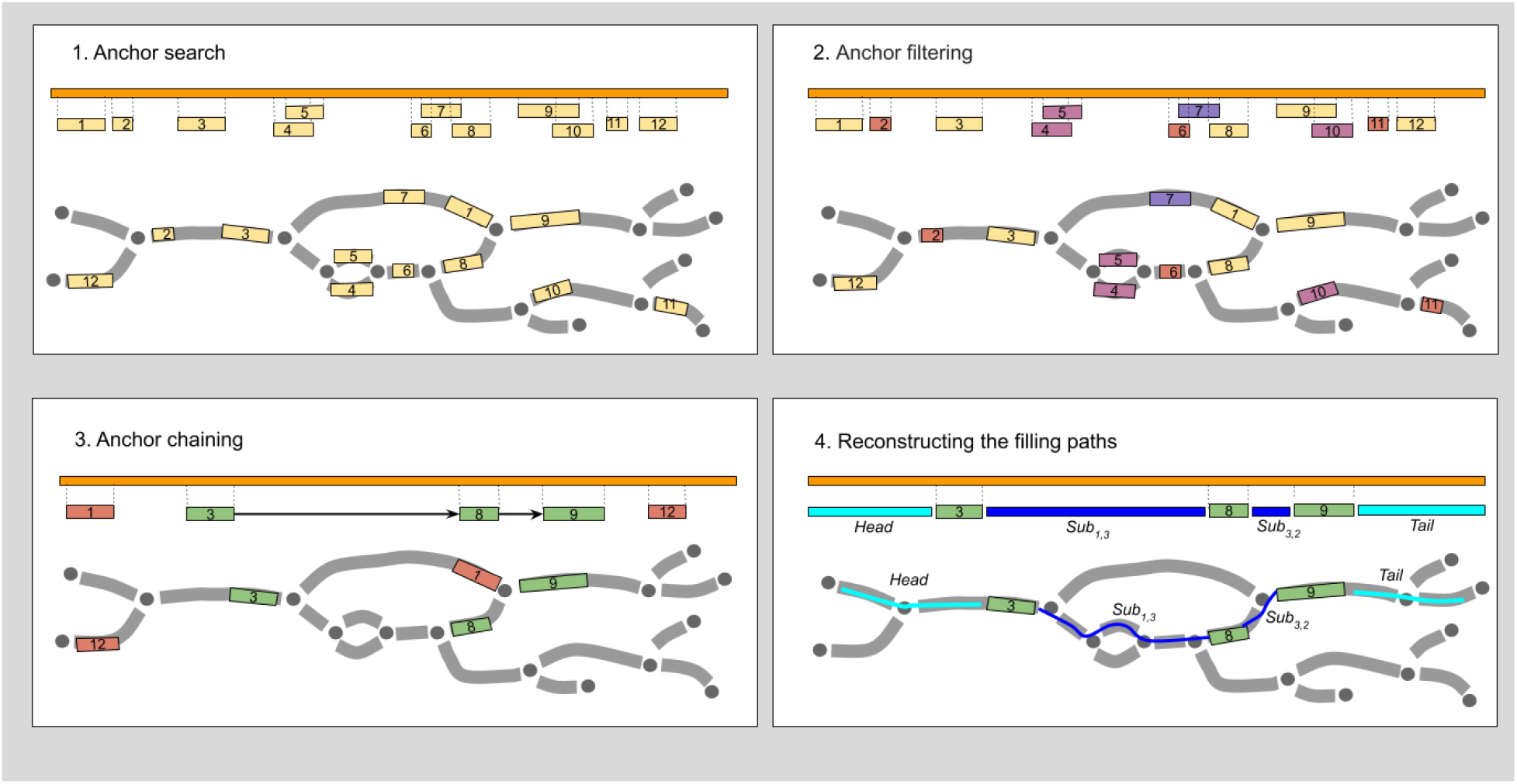
Sequence-to-graph alignment with SPAligner. Alignment of query sequence *S* (orange bar) to assembly graph *G*. Assembly graph edges are considered directed left-to-right (explicit edge orientation was omitted to improve clarity). **Upper-left: Anchor search.** Anchor alignments (regions of high similarity) between the query and the edge labels are identified with BWA-MEM. **Upper-right: Anchor filtering.** Anchors shorter than *K* (anchors 2, 6, 11), anchors “in the middle” of long edge (anchor 7) or ambiguous anchors (anchor 10 mostly covered by anchor 9, both anchors 4 and 5) are discarded. **Bottom-left: Anchor chaining.** Heaviest chain of compatible anchors (chain 3 → 8 → 9) is determined. **Bottom-right: Reconstructing the filling paths.** Paths for fragments of the query between the consecutive chain anchors (as well as left- and right-most fragments) are reconstructed.

1. **Anchor search.** Anchor alignments between *S* and the pre-indexed edge labels are identified using the BWA-MEM library [14]. Each anchor a is defined by a range on *S* — [*start*_*s*_(*a*), *end*_*s*_(*a*)) and a range on a specific edge *e*(*a*) — [*start*_*e*_(*a*), *end*_*e*_(*a*)). The span of an anchor *a* is defined as (*end*_*s*_(*a*) *start*_*s*_(*a*)). Only anchors with span exceeding a certain threshold (by default equal to *K* — the kmer size parameter of de Bruijn graph) are considered. Since the goal of this step is to get a set of all potentially-relevant anchors, BWA-MEM settings were adjusted for higher sensitivity. In particular, we modified the defaults to retain as many alignments as possible: secondary alignments were included, *mask*_*level* and *drop*_*ratio* were assigned to 20, and the seed size was decreased from 17 bp to 14 bp.
2. **Anchor filtering.** Unreliable and ambiguous anchors are further discarded based on their size, locations in the graph and the layout on the query sequence. In particular, the following groups of anchors are discarded (in the specified order):
  - Anchors “in the middle” of long edges, which span less than *T* bp (*T* = 500, by default). More formally we discard the anchor *a* if *end*_*e*_(*a*) − *start*_*e*_(*a*)+ min(*start*_*s*_(*a*), *start*_*e*_(*a*)) + *min*(|*S*| − *end*_*s*_(*a*), |*e*| − *end*_*e*_(*a*)) > 3 ⋅ (*end*_*e*_(*a*) − *start*_*e*_(*a*)).
  - Anchors for which at least half of their query ranges are covered by other anchors.
  - Anchors spanning less than *t* bp (*t* = 200, by default).
3. **Anchor chaining.** Anchors are assigned weights equal to their span. Two anchors a and b are considered compatible if the minimal distance between them in G does not significantly exceed the distance between their positions on *S*. SPAligner searches for the heaviest chain of compatible anchors using dynamic programming and considers the resulting chain as a skeleton of the final alignment [9]. Two anchor alignments *a* and *b* are considered compatible if the minimal distance between them in *G* does not significantly exceed the distance in *S* (formally, *dist*_*G*_(*p*_*a*_, *p*_*b*_)*/*(*start*_*s*_(*b*) − *end*_*s*_(*a*)) > *α*, where *p*_*a*_ = (*e*(*a*), *end*_*e*_(*a*)), *p*_*b*_ = (*e*(*b*), *start*_*e*_(*b*) and *α* defaults to 1.3).
4. **Reconstructing the filling paths.** The goal of this step is to identify paths in *G* between the consecutive anchors in the skeleton (and beyond its leftmost/rightmost anchors) minimizing the alignment cost against corresponding regions of the query sequence. Following a number of previous works on alignment of sequences against general graphs (potentially containing cycles) [15, 9, 16, 13], we use classic reformulation of the alignment problem as a minimal-weight path problem in an appropriately constructed alignment graph (or edit graph) [16, 13]. The alignment graph is a weighted directed graph which is constructed in such a way that paths between certain types of vertices unambiguously correspond that valid alignments of the query sequence to G and the alignment costs are equal to the path weights.

Though SPAligner was primarily designed for finding semi-global alignments, in some cases it may also produce split-read alignments, i.e. separate alignments covering non-overlapping segments of the query sequence (see Supplement Section “Support for split-read alignments”).

### Sequence to graph alignment via alignment graphs

While in this work we only consider linear gap penalty scoring scheme (with non-negative gap penalty parameter *σ* and mismatch cost *µ*), proposed approach can straightforwardly support affine gap cost model [17].

For the given query sequence fragment, assembly graph and the scoring parameters we define alignment graph *SG*(*G, Sub*) with vertices corresponding to all pairs of position in the graph and position in the query sequence.

The weighted ^2^ edges are introduced in *SG*(*G, Sub*) as follows:

1. *< p, pos*_*Sub*_ >→< *p*^′^, *pos*_*Sub*_ > of weight *σ*,
2. *< p, pos*_*Sub*_ >→< *p, pos*_*Sub*_ + 1 > of weight *σ*,
3. *< p, pos*_*Sub*_ >→< *p*^′^, *pos*_*Sub*_ + 1 >, of weight zero if *Sub*[*pos*_*Sub*_ + 1] matches *G*[*p*′], and *µ* otherwise,
4. *< p*_*end*_, *pos*_*Sub*_ >→< *p*_*start*_, *pos*_*Sub*_ > of weight zero if *p*_*end*_ and *p*_*start*_ both corresponds to the same vertex in *G* (i.e. *p*_*end*_ = (*e*_1_, |*e*_1_|), *p*_*start*_ = (*e*_2_, 0) and the end of edge *e*_1_ is same vertex as the start of *e*_2_),

where *p* iterates through all positions in *G*, while *p′* iterates through all graph positions extending *p* (i.e. *p* = (*e, i*) and *p′* = (*e, i* + 1) for some edge *e* and index *i* < |*e*|), and *pos*_*Sub*_ — through all (permissible) positions in *Sub*.

Consider a pair of consecutive anchors in the skeleton chain, *a*_*i*_: ([*start*_*s*_(*a*_*i*_), *end*_*s*_(*a*_*i*_)), (*e*(*a*_*i*_), [*start*_*e*_(*a*_*i*_), *end*_*e*_(*a*_*i*_)))), and *a*_*i*+1_: ([*start*_*s*_(*a*_*i*+1_), *end*_*s*_(*a*_*i*+1_), (*e*(*a*_*i*+1_), [*start*_*e*_(*a*_*i*+1_), *end*_*e*_(*a*_*i*+1_))), and a substring of query *Sub* = *S*[*end*_*s*_(*a*_*i*_): *start*_*s*_(*a*_*i*+1_)]. *ED*(*S*_1_*, S*_2_) denotes alignment cost (linear gap penalty function with parameters *µ* and *σ*) between strings *S*_1_ and *S*_2_. Our goal is to find a path *P*, connecting positions *p*_1_ = (*e*(*a*_*i*_), *end*_*e*_(*a*_*i*_)) and *p*_2_ = (*e*(*a*_*i*+1_), *start*_*e*_(*a*_*i*+1_)) in the graph *G* minimizing *ED*(*Label*(*P*), *Sub*). It is easy to see (i.e. Lemma 5 in [15]) that path *P* can be recovered from the minimal-weighted path connecting vertices *< p*_1_, 0 > and < *p*_2_, |*Sub*| > in *SG*(*G, Sub*), moreover, its weight is equal to *ED*(*Label*(*P*), *Sub*). Note that *SG*(*G, Sub*) only has non-negative weights, so minimal-weighted paths search can be efficiently performed by Dijkstra algorithm [15, 9].

In addition to reconstruction of paths between the consecutive anchors of the alignment skeleton, SPAligner also attempts to extend the alignment beyond the leftmost/rightmost anchors. Without loss of generality, we consider finding optimal alignment of the query fragment *Suf* beyond *end*_*s*_(*a*_*last*_), while fixing its starting position in the graph *G* as *p*_*s*_ = (*e*(*a*_*last*_), *end*_*e*_(*a*_*last*_)). The sought answer can be recovered from the minimal weight path in *SG*(*G, Suf*) across all paths connecting < *p*_*s*_, 0 > with any of the vertices {< *p*_*t*_, |*Suf*| >}, where *p*_*t*_ iterates through all positions in *G*, which can also be found by Dijkstra algorithm.

To speed-up the optimal path search SPAligner implements multiple heuristics (see Supplemental Section “Alignment extension heuristics and thresholds”). In particular, while we consider the entire graph *G* in the description above, a much smaller subgraph surrounding anchor alignments is considered by the SPAligner implementation. The implementation also benefits from available highly optimized solutions for sequence-to-sequence alignment (see Supplemental Text “Leveraging fast sequence-to-sequence alignment methods”).

More efficient algorithms in terms of worst case time complexity have been earlier suggested in [18, 19] for an important case of edit distance or Levenshtein distance (*µ* and *σ* equal to 1). Supplement Section “Shortest paths search in binary-weighted graphs” presents a simple modification of the basic approach described above achieving the same time complexity of *O*(|*G*| ⋅ |*Sub*|).

It worth to note that recently Jain et al.[6] suggested an elegant algorithm extending the same time complexity to both linear and affine gap penalty functions.

### Alignment of amino-acid sequences

Consider an assembly graph *G* with edges labeled by nucleotide sequences and an amino acid query sequence *S*_*p*_. We will refer to a path *P* in *G* as an *AA*-path, if its label translates into a valid (i.e.without any stop codon) protein sequence (denoted by *Translated*(*Label*(*P*))). We define semi-global alignment of a sequence *S*_*p*_ to graph *G* as an *AA*-path *P* in *G*, such that the alignment cost between *S*_*p*_ and *Translated*(*Label*(*P*)) is minimal across all *AA*-paths in *G* ^3^.

While considering fixed mismatch penalties is usually sufficient for aligning nucleotide sequences, most practical scoring schemes for amino acid sequence alignment use specialized substitution matrices (e.g. BLOSUM or PAM matrices) [20], which take into account the substitution rates between different pairs of amino acids over time as well as the relative frequencies of various amino acids.

To reconcile our cost-minimization formulation with the fact that typical substitution matrix *M* assumes better alignments to have higher cumulative scores, we will consider elements of matrix *M′* = *M* as match/mismatch score (by default *M* =BLOSUM90).

Again, while we only consider linear gap penalties with coefficient *σ* (in our experiments we used *σ* = 5) here, affine gap penalties can also be implemented without significant increase of running time or memory footprint [19].

### Alignment graph for amino acid sequence alignment

For the given query sequence, assembly graph, substitution matrix and gap penalty coefficient (*σ*), we introduce alignment graph, *SG*_*p*_. Each vertex corresponds to a triplet 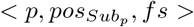, where *p* is position in the graph *G*, 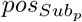 is the position in the query fragment sequence and *f s* is a frame state string (of length 0 − 2), encoding a partial codon sequence (empty string is denoted by *ε*). We further define the graph by the iterative procedure, initialized by introducing the vertices (*p*, 0, *ε*), where *p* iterates through all positions in *G*. Then, until no more edge can be added, we introduce edges and required vertices following one of the rules below:

1. 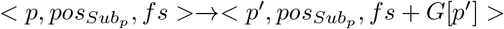 of zero weight if *|f s| <* 2,
2. 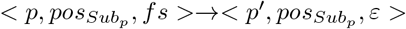 of weight *σ* if *|f s|* = 2,
3. 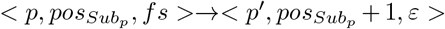 of weight 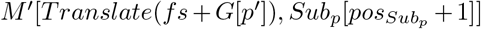, and *|f s|* = 2,
4. 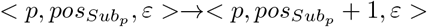 of weight *σ*,
5. 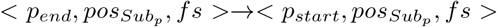 of weight zero if *p*_*end*_ and *p*_*start*_ correspond to the same position in graph on consecutive edges of *G* (i.e. *p*_*end*_ = (*e*_1_, |*e*_1_|), *p*_*start*_ = (*e*_2_, 0) and end of edge *e*_1_ is same vertex as the start of *e*_2_),

where *p* is a position in *G*, *p′* is a position in *G* extending *p* (i.e. *p* = (*e, i*) and *p′* = (*e, i* + 1) for some edge *e* and index *i* < |*e*|),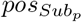 – is a position in *Sub*_*p*_.

As before, one can find the semi-global alignment path of the sequence *Sub*_*p*_ (fixed on one or both sides by anchor alignments) by searching for minimal weight paths in *SG*_*p*_.

In particular, it is easy to show that the semi-global alignment of the amino acid sequence *Sub*_*p*_ starting from position *p*_*s*_ in the graph *G* corresponds to the minimal weight path in *SG*_*p*_(*G, Sub*_*p*_) connecting vertex < *p*_*s*_, 0, *ε* > to any of the vertices {< *p*_*t*_, |*Sub*_*p*_|, 0 >}, where *p*_*t*_ iterates through all positions in *G*.

Note that since any reasonable matrix *M′* contains negative elements, the graph *SG*_*p*_ will have edges of negative weight, preventing direct application of Dijkstra algorithm for the search of such minimal weight paths. While Algorithm 2 from [19] (slightly modified to account for the frame states) can be applied here, we use an alternative (and more straightforward) approach resulting in the same time complexity.

We define a new graph 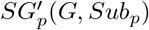 increasing the weights of edges introduced by rules 3 and 4 by an arbitrary constant *C*, exceeding the maximum value in *M*. Since the resulting graph has only edges with non-negative weights, Dijkstra algorithm can be used for the shortest paths search in this graph.

At the same time it is easy to show that the minimum weight path between two arbitrary nodes of the form *s* =< *x*, 0, *ε* > and *e* =< *y*, |*Sub*_*p*_| + 1, *ε* > in 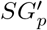 is also the minimum weight path between *s* and *e* in *SG*_*p*_. Indeed, if we consider an arbitrary path *P* in *SG*_*p*_ between *s* and *e* of weight *w*_*P*_, its weight after transformation (in 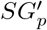) is *w*_*p*_ + *C* ⋅ |*Sub*_*p*_|, because *P* contains exactly |*Sub*_*p*_| edges with modified weights. Therefore, the weights of all paths between two vertices *s* and *e* will be increased by the same constant |*Sub*_*p*_| ⋅ *C*.

### Amino acid sequence alignment pipeline

Below we outline the four steps implemented in SPAligner for aligning an amino acid sequence *S*_*p*_ to an assembly graph *G*, highlighting the differences with the pipeline presented in section “Alignment of long nucleotide sequences”.

1. **Anchors search.** As before SPAligner first identifies anchor alignments between *S*_*p*_ and the pre-indexed six-frame translated edge labels. Current implementation relies on finding high-scoring local alignments between the “canonical” nucleotide representations of protein sequences, obtained by substituting every amino acid by its lexicographically minimal codon. The search is performed by BWA-MEM configured to prevent gapped alignments (*gap*_*open* penalty set to infinity) and we additionally check that the aligned ranges are consistently located with respect to the reading frame.
2. **Anchor filtering.** Unreliable and ambiguous anchors are discarded based on their size and query range. Anchors shorter than K are filtered out. Smaller lengths of typical protein sequences as compared to long read sequences allow us to omit the anchor chaining step in order to increase the accuracy of the resulting amino acid sequence alignments. It also eliminates the need to coordinate the reading frames of the chained anchors.
3. **Alignment extension.** SPAligher then uses the adjusted alignment graph approach (see section “Alignment graph for amino acid sequence alignment”) to search for optimal alignments extending each of the remaining anchors. The procedure employs several heuristics and constraints, which can be found in Supplemental Section “Alignment extension heuristics and thresholds”.
4. **Alignments post-processing.** SPAligner removes alignment paths shorter than a certain fraction of the query coding sequence length (0.8 by default) and filters perfectly duplicating paths. Resulting paths are then translated and re-scored against the query sequence with Parasail library [21].

## Results and discussion

### Aligning long error-prone reads

We benchmarked SPAligner against the two sequence-to-graph aligners supporting long reads – vg *v*1.17 and GraphAligner *v*1.0.4.

The vg alignment pipeline [4] starts from searching for super maximal equal matches (SMEMs) with GCSA2 library [22]. After filtering and chaining of SMEMs, the SMEMs in a chain are linked by SIMD-accelerated banded dynamic programming. In order to use this method, vg first “unrolls” the cycles to transform the graph into a directed acyclic graph (DAG). It is worth noting that this step can produce large intermediate graphs and potentially lead to suboptimal alignments. While working with long sequences, vg splits them into overlapping “chunks” (default 256 bp with 32 bp overlap) and maps them separately, further combining them by the same method as for SMEMs chaining.

GraphAligner [13] uses MUMmer4 [23] to identify potential alignment seeds to restrict the search for the potential alignment paths. Each seed is then extended separately by a novel bitvector-based algorithm for semi-global sequence-to-graph alignment under an edit distance (unit costs) model. Based on simultaneous processing of *w* sequence characters, where *w* is the word size of the machine and presented as a generalization of Myers’ bit-vector algorithm [24], it is reported to achieve considerable practical speedup over previously suggested asymptotically optimal algorithms [18, 13].

Besides the default configuration of GraphAligner we have also benchmarked its “try-all-seeds” mode (disabling the filtering of likely false positive seed alignments), recommended by the developers for bacterial datasets.

For benchmarking we considered PacBio and ONT sequencing reads for three different organisms: *E. coli* strain K12, *S. cerevisiae* strain S288C and *C. elegans* strain Bristol N2 (see Supplemental Text “Datasets availability” for accession numbers). Assembly graphs were generated with SPAdes *v*3.12 [1] assembler from the appropriate short-read Illumina datasets (using “-k 21,33,55,77” and setting other parameters to defaults). While running the aligners we tried to follow the recommendations given by the developers (see Supplemental Text “Notes on running aligners” for details). Benchmarking results are summarized in Table 1, with columns corresponding to the following metrics:

**Table 1:** Summary statistics of aligning PacBio/ONT reads to short-read assembly graphs (constructed by SPAdes with kmer size equal to 77). Every dataset consists of 10k reads longer than 2Kbp (except *E. coli* ONT dataset with 7k reads of appropriate length). All runs performed in 16 threads. Due to high fragmentation of assembly graph resulting from available *C. elegans* Bristol N2 Illumina sequencing data, we used simulated Illumina reads to obtain *C. elegans* assembly graph.

| Aligner | MR(%) | AI (%) | T (days<br>h:m) | M (Gb) |  | MR(%) | AI (%) | T (days<br>h:m) | M (Gb) |
| --- | --- | --- | --- | --- | --- | --- | --- | --- | --- |
| *E. coli* PacBio |  |  |  |  |  | *E. coli* ONT |  |  |  |
| vg | 29 | 90 | 00:41 | 4.1 |  | 58 | 88 | 00:33 | 3.3 |
| GraphAligner | 85 | 88 | 00:01 | 0.1 |  | 82 | 87 | 00:01 | 0.2 |
| GA try-all-seeds | 86 | 88 | 00:25 | 0.3 |  | 84 | 87 | 01:43 | 4.8 |
| SPAligner | 88 | 87 | 00:04 | 0.3 |  | 86 | 87 | 00:03 | 0.3 |
| *S. cerevisiae* PacBio |  |  |  |  |  | *S. cerevisiae* ONT |  |  |  |
| vg | 15 | 90 | 1 day<br>04:03 | 91 |  | 70 | 47 | 42:59 | 85 |
| GraphAligner | 53 | 87 | 00:01 | 0.2 |  | 65 | 82 | 00:01 | 0.2 |
| SPAligner | 56 | 87 | 00:38 | 0.7 |  | 61 | 83 | 00:16 | 0.6 |
| *C. elegans* PacBio |  |  |  |  |  | *C. elegans* ONT |  |  |  |
| vg | 10 | 91 | 3 days<br>04:26 | 306 |  | 15 | 89 | 2 days<br>12:58 | 309 |
| GraphAligner | 84 | 88 | 00:03 | 1.8 |  | 63 | 87 | 00:03 | 1.8 |
| SPAligner | 84 | 87 | 01:06 | 1.2 |  | 65 | 87 | 00:34 | 1.3 |

- *Number of mapped reads* (MR): Read is considered mapped if 80% of its length is covered by a single alignment path.
- *Alignment identity (%)*(AI): Mean nucleotide identity ^4^ across the longest continuous alignments of the mapped reads.
- *Wall-clock time (days h:m)*(T): Wall-clock time with 16 threads.
- *Memory (Gb)*(M): Peak RAM usage with 16 threads.

Vg showed highest average alignment identity for all considered datasets, but at the same time resulted in very low fraction of mapped reads. The same tendency can be seen on synthetic datasets (see Supplementary Table S2).

SPAligner showed 2-4% advantage in number of mapped reads over GraphAligner for *E. coli* datasets, while GraphAligner was able to map 4% more reads from *S. cerevisiae* ONT dataset. It is worth to note that the size of the intersection of reads mapped by SPAligner and GraphAligner (equal to 84% for *E. coli* PacBio dataset and 60% for *S. cerevisiae* ONT) is close to the number of reads mapped by each tool and average identity of SPAligner and GraphAligner on the set of reads mapped by both tools is the same — 88% and 83% for *E. coli* and *S. cerevisiae* respectively (see Supplementary Table S3). Thus we can conclude that in both cases differences were caused by successful alignments of additional reads with lower identity. At the same time SPAligner was able to successfully reconstruct more alignments on E. coli ONT data saving the same identity level — 87%.

Results of additional benchmarking on the synthetic PacBio and ONT datasets are summarized in Supplementary Table S2.

Overall, our benchmarking suggests that SPAligner and GraphAligner produce similar alignments of long error-prone reads with SPAligner being faster and more memory-efficient than GraphAligner in “try-all-seeds” mode and slower and less memory-efficient than GraphAligner in default mode.

### Aligning protein sequences

In contrast to other sequence-to-graph aligners available at the moment, SPAligner supports alignment of protein sequences onto assembly graphs. In this section we demonstrate how SPAligner can be efficiently used to identify regions of the assembly graph exhibiting high similarity to known prokaryotic protein sequences.

Contigs originating from metagenomic assemblies can be excessively fragmented due to various reasons, including inter-species repeats (conservative sequences, horizontally transferred genes, etc) and within-species microdiversity [25]. Contig fragmentation can lead to loss of important insights in modern metagenomic studies, which increasingly rely on annotation of assembled contigs for functional analysis of the microbial community members [26, 27]. Alignment of known protein sequences onto metagenomic assembly graph provides a complementary approach capable of identifying genes fragmented over several contigs.

The capacity of applying SPAligner in this context is demonstrated below in two experiments, considering alignments of:

- coding sequences from entire UniProt (The UniProt Consortium, 2019) database onto an assembly graph of a well-defined synthetic metagenomic community;
- a comprehensive set of known antibiotic resistance genes onto an assembly graph for a wastewater metagenomic dataset.

While alignment of a protein sequence with SPAligner might result in several alignment paths (initiated by different anchors), in this work we will only consider single best scoring alignment for each query.

### Annotating genes in synthetic metagenomic community

Shakya et al. [28] performed sequencing of synthetic DNA mixture for 64 diverse bacterial and archaeal organisms (SRA acc. no. SRX200676). This dataset, containing 109 million 2 *×* 100 bp Illumina reads (with mean insert size of 206 bp) has been extensively used for benchmarking of tools for metagenomic analysis [29, 30]. Reference genome annotations available for all organisms forming the community contain a total of 196150 protein coding sequences, further referred as reference protein sequences. We model the situation in which we aim to identify potential close homologs of known protein sequences within the metagenomic assembly graph. In particular, we searched for semi-global alignments of 350 thousand sequences comprising “reviewed” UniProt database of bacterial proteins [31]. In order to identify UniProt sequences that are expected to align to the assembly graph (assuming that the graph represents all necessary sequences), we aligned UniProt sequences to the set of reference protein sequences. We used BLASTp [32] with default settings in order to quickly find most similar sequences to the reference one. As a result, 26209 UniProt sequences were mapped to some reference protein with 90% amino acid identity threshold. We consider assembly graph, built by metaSPAdes *v*3.12 [29] with default settings, and use SPAligner to identify alignments of UniProt sequences at 90% amino-acid identity threshold. SPAligner was able to align 93% (24324 out of 26209) of UniProt sequences with matching reference proteins (see above), as well as 1299 extra sequences. Further analysis showed that the majority of falsely identified sequences are similar to substrings of some reference protein sequences.

### Identification of antibiotic resistance genes in a hospital wastewater sample

In this section we demonstrate how SPAligner can be applied to identify genes of particular interest within real metagenomic dataset.

Spread of antimicrobial resistance is an escalating problem and a threat to public health. Metagenomic sequencing provides an efficient methodology for detection and tracking of the antibiotic resistance genes (ARGs) in the environment samples. Ng et al. [33] explored wastewater and urban surface water metagenomic datasets for the presence of various antimicrobial resistance (AMR) proteins, in particular beta-lactamases. To illustrate the ability of SPAligner to identify additional AMR gene families as compared to conventional approaches based on analysis of assembled contigs, we focus on a hospital wastewater discharge dataset, which, according to the original study, had the highest total coverage of beta-lactamase genes. The dataset consisting of 3.3 million 2 × 250 bp Illumina reads with mean insert size of 350 bp (SRA acc. no. SRR5997548) was assembled by metaSPAdes *v*3.12 with default settings.

First, we used state-of-the-art AMRFinder tool [34] to identify AMR genes in assembled contigs. AMRFinder performs the BLASTx [32] searches of the protein sequences from Bacterial Antimicrobial Resistance Reference Gene Database [34], containing protein sequences for over 800 gene families representing 34 classes of antimicrobials and disinfectants. Since we are aiming at the identification of the (almost) complete protein sequences, partial predictions covering no more than 75% of the corresponding database protein were discarded. All other parameters were left to defaults, including the 90% alignment identity threshold.

We then used SPAligner to identify best alignments (with at least 90% amino-acid identity threshold) for all 4810 ARG sequences from the same database onto the graph.

To check whether our sequence-to-graph alignments were able to capture additional AMR protein families as compared to the baseline contig analysis approach, protein sequences identified by AMRFinder and SPAligner were clustered by single-linkage clustering with a 90% similarity cut-off. Clusters where only SPAligner alignments are presented tend to be really far by identity from alignments found by AMRFinder and vice versa. Out of the resulting 89 clusters a single cluster consisted of only AMRFinder prediction. Further analysis showed that SPAligner also successfully identified corresponding protein sequence which spanned 89.7% of the query length, while AMRFinder prediction spans 90.7%. At the same time, 4 clusters contained only SPAligner’s alignments. tBLASTx [32] alignments of the corresponding database protein sequences onto the assembled contigs either covered less than 75% of the query or had low identity (*<* 70%).

Figure 2 illustrates the alignment paths of beta-lactamase sequences IMP-1 and IMP-4, forming one of the 4 clusters with exclusively SPAligner-based predictions. For each of those genes SPAligner had identified an alignment path distributed across several contigs covering the entire query sequence with *>* 98% identity.

**Figure 2:**
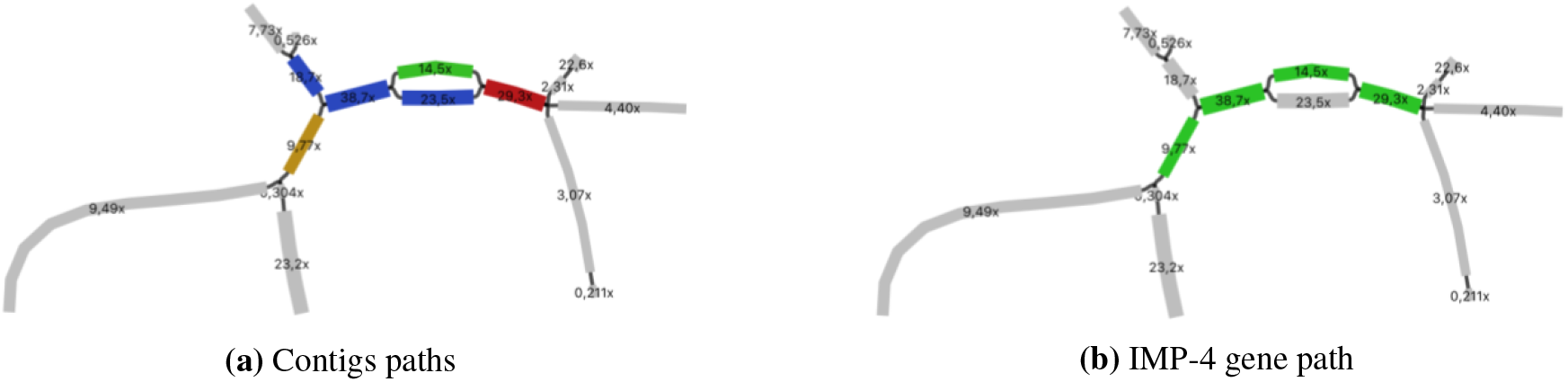
Example of putative beta-lactamase protein sequence, fragmented in the metagenomic assembly contigs. a) Relevant region of assembly graph visualized by Bandage software [35]. Blue, red, green and brown colors correspond to 4 different contigs within the metaSPAdes assembly. b) Visualization of the alignement path for IMP-4 gene (246 aa long, alignment identity — 98%). Edges comprising the path are colored in green.

We further relate our analysis to the original study [33] with respect to the identified beta-lactamase families. Alignments were obtained for instances of seven out of nine beta-lactamase protein families from the original study (*bla*_*KPC*_, *bla*_*CTX−M*_, *bla*_*SHV*_, *bla*_*TEM*_, *bla*_*IMP*_, *bla*_*VIM*_ and *bla*_*OXA*_). While SPAligner was not able to produce alignments for two remaining gene families (*bla*_*NDM*_ and *bla*_*CMY*_), in the original study their instances were estimated to be poorly covered compared to other families ^5^.

Note that in [33] authors tried to identify multiple beta-lactamase sequences within the same protein family, which we are not aiming to do here, since minor differences within highly similar gene sequences are likely to be lost in the metagenomic assembly process. Such high-resolution analysis should be possible within the sequence-to-graph alignment framework via configuring the assembly graph construction stage to preserve necessary genomic variation. This promising direction is left for future research.

### Future improvements

In particular, our results suggest that SPAligner is an efficient solution for mapping third generation sequencing data and can also facilitate the identification of known genes in complex metagenomic datasets.

Besides general improvements of the codebase, we are planning to migrate to more efficient libraries for searching anchor alignments [23, 22] and implement the more efficient extension algorithms [6, 13].

An important direction with respect to the nucleotide sequence alignment is improving the alignment of reads structurally different from the genome, represented by the assembly graph [36]. This includes a more reliable strategies for ignoring poorly sequenced read extremities and producing split-read alignments.

With respect to the amino acid sequence alignment our future goals include improving the sensitivity of the anchor alignment to allow for searching of the more diverged protein sequences, as well as development of strategies for identification of multiple closely-related gene sequences within the custom metagenomic assembly graphs with preserved variation.

## Conclusions

In this work we present SPAligner tool for aligning long diverged molecular sequences to assembly graphs and demonstrated its effectiveness for aligning both reads produced by third-generation sequencing technologies and protein sequences.

SPAligner was benchmarked against the two sequence-to-graph aligners supporting long reads – vg and GraphAligner, and showed highly competitive results on various Pacbio and Oxford Nanopore datasets. At the same time being the only tool that produce alignment of amino acid sequences, SPAligner was successfully used to identify gene sequences, which were not fully recovered by metagenomic assembly.

## Availability and requirements

**Project name:** SPAligner

**Project home page:** https://github.com/ablab/spades/archive/spaligner-paper.zip

**Operating system(s):** Platform independent

**Programming language:** C++

**Other requirements:** g++ (version 5.3.1 or higher), cmake (version 2.8.12 or higher), zlib, libbz2

**License:** GNU GPLv2

**Any restrictions to use by non-academics:** None

## List of abbreviations

ED: edit distance
ARG: antibiotic resistance gene
AMR: antimicrobial resistance

## Declarations

### Ethics approval and consent to participate

Not applicable.

### Consent for publication

Not applicable.

### Availability of data and materials

Source code and manual of SPAligner tool is available on https://github.com/ablab/spades/archive/spaligner-paper.zip. The datasets analysed during the current study are available in https://figshare.com/s/b0dc3f715e0224e3e962.

### Authors’ contributions

SN suggested the idea. TD is the main author of SPAligner implementation. DA took part in implementation of anchor chaining part. AK integrated BWA-MEM module. TD conducted SPAligner comparison with other tools and interpreted the data. SN, DA and AK revised the tool algorithms. SN, TD and DA prepared the manuscript. All authors read and approved the final version of the manuscript.

## Acknowledgements

We wish to thank Alexander Shlemov for great ideas and all the members of Center for Algorithmic Biotechnology for the helpful discussion.

## Footnotes

1 While common definition of assembly graph associates sequences with the vertices, assembly graphs produced by de Bruijn graph based assemblers can be readily represented as edge-labeled graphs, considered in this work. We also note that our methods can be straightforwardly re-implemented in application to other assembly graphs.

2 We use “weights” rather than “lengths” notation here to simplify introduction of negative weights in later sections.

3 Note that here we are not interested in paths spelling the “frame-shifted” version of the protein coding sequence (in particular, |*Label*(*P*)| is always divided by 3). Here we implicitly rely on the assumption that the assembly graph was constructed from highly accurate reads and thus rare “frame shift” events need not be considered.

4 Identity of aligning sequence *S* onto path *P* is defined as 1 *− ED*(*S, Label*(*P*))/|*S*|.

5 Total abundance of the identified families exceeds 150X, while the total coverage of NDM and CMY families was estimated as 20X. The exact formula for abundance calculation is presented in the original study [33].

## References

[1] Sergey Nurk, Anton Bankevich, Dmitry Antipov, Alexey Gurevich, Anton Korobeynikov, Alla Lapidus, Andrey Prjibelsky, Alexey Pyshkin, Alexander Sirotkin, Yakov Sirotkin, Ramunas Stepanauskas, Jeffrey McLean, Roger Lasken, Scott R. Clingenpeel, Tanja Woyke, Glenn Tesler, Max A. Alekseyev, and Pavel A. Pevzner. Assembling genomes and mini-metagenomes from highly chimeric reads. In Minghua Deng, Rui Jiang, Fengzhu Sun, and Xuegong Zhang, editors, Research in Computational Molecular Biology, volume 7821, pages 158–170. Springer Berlin Heidelberg.

[2] Rayan Chikhi and Guillaume Rizk. Space-efficient and exact de bruijn graph representation based on a bloom filter. In WABI, volume 7534 of Lecture Notes in Computer Science, pages 236–248. Springer.

[3] Dinghua Li, Chi-Man Liu, Ruibang Luo, Kunihiko Sadakane, and Tak-Wah Lam. MEGAHIT: an ultra-fast single-node solution for large and complex metagenomics assembly via succinct de bruijn graph. 31(10):1674–1676.

[4] Erik Garrison, Jouni Sirén, Adam M Novak, Glenn Hickey, Jordan M Eizenga, Eric T Dawson, William Jones, Shilpa Garg, Charles Markello, Michael F Lin, Benedict Paten, and Richard Durbin. Variation graph toolkit improves read mapping by representing genetic variation in the reference. 36:875.

[5] Mahdi Heydari, Giles Miclotte, Yves Van de Peer, and Jan Fostier. BrownieAligner: accurate alignment of illumina sequencing data to de bruijn graphs. 19(1):311.

[6] Chirag Jain, Haowen Zhang, Yu Gao, and Srinivas Aluru. On the complexity of sequence to graph alignment.

[7] Vaddadi Naga Sai Kavya, Kshitij Tayal, Rajgopal Srinivasan, and Naveen Sivadasan. Sequence alignment on directed graphs.

[8] Antoine Limasset, Bastien Cazaux, Eric Rivals, and Pierre Peterlongo. Read mapping on de bruijn graphs. 17(1):237–237.

[9] Dmitry Antipov, Anton Korobeynikov, Jeffrey S. McLean, and Pavel A. Pevzner. hybridSPAdes: an algorithm for hybrid assembly of short and long reads. 32(7):1009–1015.

[10] Ryan R. Wick, Louise M. Judd, Claire L. Gorrie, and Kathryn E. Holt. Unicycler: Resolving bacterial genome assemblies from short and long sequencing reads. 13(6):e1005595.

[11] Leena Salmela and Eric Rivals. LoRDEC: accurate and efficient long read error correction. 30(24):3506–3514.

[12] Shilpa Garg, Mikko Rautiainen, Adam M Novak, Erik Garrison, Richard Durbin, and Tobias Marschall. A graph-based approach to diploid genome assembly. 34(13):i105–i114.

[13] Mikko Rautiainen, Veli Mäkinen, and Tobias Marschall. Bit-parallel sequence-to-graph alignment.

[14] Heng Li. Aligning sequence reads, clone sequences and assembly contigs with BWA-MEM.

[15] Amihood Amir, Moshe Lewenstein, and Noa Lewenstein. Pattern matching in hypertext. 35(1):82–99.

[16] Eugene W. Myers. AnO(ND) difference algorithm and its variations. 1(1):251–266.

[17] Osamu Gotoh. An improved algorithm for matching biological sequences. 162(3):705–708.

[18] Gonzalo Navarro. A guided tour to approximate string matching. 33(1):31–88.

[19] Mikko Rautiainen and Tobias Marschall. Aligning sequences to general graphs in (+) time.

[20] William R. Pearson. Selecting the right similarity-scoring matrix: Selecting the right similarity-scoring matrix. In Alex Bateman, William R. Pearson, Lincoln D. Stein, Gary D. Stormo, and John R. Yates, editors, Current Protocols in Bioinformatics, pages 3.5.1–3.5.9. John Wiley & Sons, Inc.

[21] Jeff Daily. Parasail: SIMD c library for global, semi-global, and local pairwise sequence alignments. 17(1):81.

[22] Jouni Sirén. Indexing variation graphs. pages 13–27.

[23] Guillaume Marçais, Arthur L. Delcher, Adam M. Phillippy, Rachel Coston, Steven L. Salzberg, and Aleksey Zimin. MUMmer4: A fast and versatile genome alignment system. 14(1):e1005944.

[24] Gene Myers. A fast bit-vector algorithm for approximate string matching based on dynamic programming. In Martin Farach-Colton, editor, Combinatorial Pattern Matching, volume 1448, pages 1–13. Springer Berlin Heidelberg.

[25] Niranjan Nagarajan and Mihai Pop. Sequence assembly demystified. 14(3):157–167.

[26] Tyler P. Barnum, Israel A. Figueroa, Charlotte I. Carlström, Lauren N. Lucas, Anna L. Engelbrektson, and John D. Coates. Genome-resolved metagenomics identifies genetic mobility, metabolic interactions, and unexpected diversity in perchlorate-reducing communities. 12(6):1568–1581.

[27] Itai Sharon, Michael Kertesz, Laura A. Hug, Dmitry Pushkarev, Timothy A. Blauwkamp, Cindy J. Castelle, Mojgan Amirebrahimi, Brian C. Thomas, David Burstein, Susannah G. Tringe, Kenneth H. Williams, and Jillian F. Banfield. Accurate, multi-kb reads resolve complex populations and detect rare microorganisms. 25(4):534–543.

[28] Migun Shakya, Christopher Quince, James H. Campbell, Zamin K. Yang, Christopher W. Schadt, and Mircea Podar. Comparative metagenomic and rRNA microbial diversity characterization using archaeal and bacterial synthetic communities: Metagenomic and rRNA diversity characterization. 15(6):1882–1899.

[29] Sergey Nurk, Dmitry Meleshko, Anton Korobeynikov, and Pavel A. Pevzner. metaSPAdes: a new versatile metagenomic assembler. 27(5):824–834.

[30] Sherine Awad, Luiz Irber, and C. Titus Brown. Evaluating metagenome assembly on a simple defined community with many strain variants.

[31] A. Bairoch. The SWISS-PROT protein sequence database and its supplement TrEMBL in 2000. 28(1):45–48.

[32] Stephen F. Altschul, Warren Gish, Webb Miller, Eugene W. Myers, and David J. Lipman. Basic local alignment search tool. 215(3):403–410.

[33] Charmaine Ng, Martin Tay, Boonfei Tan, Thai-Hoang Le, Laurence Haller, Hongjie Chen, Tse H. Koh, Timothy M. S. Barkham, Janelle R. Thompson, and Karina Y.-H. Gin. Characterization of metagenomes in urban aquatic compartments reveals high prevalence of clinically relevant antibiotic resistance genes in wastewaters. 8.

[34] Michael Feldgarden, Vyacheslav Brover, Daniel H. Haft, Arjun B. Prasad, Douglas J. Slotta, Igor Tolstoy, Gregory H. Tyson, Shaohua Zhao, Chih-Hao Hsu, Patrick F. McDermott, Daniel A. Tadesse, Cesar Morales, Mustafa Simmons, Glenn Tillman, Jamie Wasilenko, Jason P. Folster, and William Klimke. Using the NCBI AMRFinder tool to determine antimicrobial resistance genotype-phenotype correlations within a collection of NARMS isolates.

[35] Ryan R. Wick, Mark B. Schultz, Justin Zobel, and Kathryn E. Holt. Bandage: interactive visualization of de novo genome assemblies: Fig. 1. 31(20):3350–3352.

[36] Fritz J. Sedlazeck, Philipp Rescheneder, Moritz Smolka, Han Fang, Maria Nattestad, Arndt von Haeseler, and Michael C. Schatz. Accurate detection of complex structural variations using single-molecule sequencing. 15(6):461–468.

