## Supplementary File for "SPAligner: Alignment of Long Diverged Molecular Sequences to Assembly Graphs"

#### Supplementary Material for "SPAligner: alignment of long diverged molecular sequences to assembly graphs"

### 1 Alignment extension heuristics and thresholds

In this section we list the heuristics and constraints used by SPAligner during the alignment extension process (e.g. construction and processing of the alignment graphs). Despite significantly improving the overall running-time, some of them may also result in recovery of sub-optimal alignment paths. The default parameter values were set to maintain reasonable speed/accuracy trade-offs.

- While constructing an alignment graph $SG(G, S)$ for finding an optimal alignment path of nucleotide sequence $S$ between positions $s$ and $e$ in graph $G$, only the subgraph of $G$ representing the union of paths shorter than $\alpha \cdot |S|$ ($\alpha = 1.3$ by default) between $s$ and $e$ is considered.
- Search for the optimal filling path between consecutive anchors in the alignment skeleton (see section "Sequence to graph alignment via alignment graphs") is aborted if number of vertices in the corresponding alignment graph exceeds $max\_gs\_states$ threshold (by default equal to $120 \cdot 10^6$).
- Maximal weight of the optimal path in the alignment graph $SG(G, S)$ is limited by a fraction of the length of nucleotide sequence $|S|$ (by default $|S|/5$).
- While searching for an optimal alignment path of nucleotide sequence $S$ between positions $s$ and $e$ in graph $G$ a straightforward enumeration of a limited number of paths between the two positions (by default 5000) is performed. Edlib library [1] is used to find the alignment scores of their sequences against $S$. Minimal attained score is then used to further bound the weight of the optimal path.
- While finding the optimal alignment paths of the nucleotide sequence fragment $Suf$ beyond rightmost anchor, the minimal alignment score $min\_score(i)$ is maintained for every prefix $Suf[0 : i]$. At any moment, the score for any alignment graph vertex, corresponding to the prefix $Suf[0 : i]$ is additionally bounded by $min\_score(i) + i \cdot penalty\_ratio$ ($penalty\_ratio = 0.1$ by default). To further prevent potential performance issues, the search is performed only if $|Suf|$ does not exceed a certain threshold (5 Kb by default).
- Upper bounds are introduced on the number of entries within the priority queue within Dijkstra algorithm as well as the total number of queue extraction events (default value is $10^6$ for both). Whenever any of the limits is exceeded, the search is aborted.
- While aligning amino acid sequences, the alignment path can not be extended beyond a stop codon.

### 2 Support for split-read alignments

SPAligner was primarily developed to find semi-global sequence alignments. But if the path satisfying the constraints for the entire query can not be identified, the procedure might result in several (query-disjoint) alignments. For example, whenever the search for an appropriate path between two consecutive skeleton anchors fails (see Section "Alignment of long nucleotide sequences"), the alignment is divided into two parts.

Moreover, in the general case, anchor chaining step is repeated several times for a single query. After the skeleton chain is identified anchors with query ranges spanned by the chain are discarded. The process is repeated while there are anchor alignments to consider.

Reconstruction of the filling paths is then invoked independently for each resulting chain.

### 3 Shortest paths search in binary-weighted graphs

Straightforward approach of aligning query $S$ to graph $G$ (both with fixed or arbitrary start/end positions) via searching the appropriate minimal-weight path in alignment graph with Dijkstra algorithm (see section "Sequence to graph alignment via alignment graphs"; [2, 3]) has the worst-case time complexity of $O(|G| \cdot |S| \cdot \log(|G| \cdot |S|))$. Several improved algorithms [4, 5] with the time complexity of $O(|G| \cdot |Sub|)$ have been earlier suggested for an important case of $\mu$ and $\sigma$ equal to 1 (edit distance alignment cost).

We note that the same time complexity can be achieved by a slight modification of Dijkstra algorithm.

The edges of $SG(G, Sub)$ under the edit distance score have weights of either 0 or 1. It is easy to see that at any moment of searching for the shortest path in such binary-weighted graph, the priority queue within Dijkstra algorithm contains elements with no more than two distinct priority values, which can only differ by 1. Thus the priority queue can be replaced by a deque: all the vertices across 0-edges (1-edges) are appended along with the corresponding distance to the beginning (to the end) of deque. Deque implementation provides all necessary operations in $O(1)$ time, improving the overall running time to $O(|G| \cdot |S|)$.

#### 4 Leveraging fast sequence-to-sequence alignment methods

We implemented a modification of the approach described in Section "Sequence to graph alignment via alignment graphs", which in practice benefits from available highly optimized solutions for sequence alignment. We define a new graph $SG(G, Sub)$ over a subset of vertices of $SG(G, Sub)$. For two particular anchors $a$ and $b$ $SG(G, Sub)$ have vertices corresponding to

- $< p_a, 0 >$, where $p_a = (e(a), end_e(a))$,
- $< p_b, |Sub| >$, where $p_b = (e(b), start_e(b))$,
- $< p_{start}(e), pos_{Sub} >$ and $< p_{end}(e), pos_{Sub} >$, where $p_{start}(e) = (e, 0)$, $p_{end}(e) = (e, |e|)$ and $e$ iterates through all edges of $G$ and $pos_{Sub} \in [0, |Sub|]$.

Edges of $SG(G, Sub)$ are defined as follows:

- $< p_{start}(e), pos_{Sub} > \to < p_{end}(e), pos_{Sub} + c >$ of length $ED(e, Sub[pos_{Sub} : pos_{Sub} + c])$,
- $< p_a, 0 > \to < p_end(e(a)), c >$ of length $ED(e(a)[end_e(a) : |e(a)|], Sub[0 : c])$,
- $< p_start(e(b)), pos_Sub > \to < p_b, |Sub| >$ of length $ED(e(b)[0 : start_e(b)], Sub[pos_{Sub} : |Sub|]$,
- $< p_end, pos_Sub > \to < p_start, pos_Sub >$ of weight zero if $p_{end}$ and $p_{start}$ both corresponds to the same vertex in $G$ (i.e. $p_{end} = (e_1, |e_1|)$, $p_{start} = (e_2, 0)$ and the end of edge $e_1$ is same vertex as the start of $e_2$),

where $e$ iterates through all edges of $G$ and $Sub_{pos}$ and $c$ – through all permissible positions in $Sub$.

While searching for the shortest path in $SG(G, Sub)$, with Dijkstra algorithm, SPAligner calculates distances from vertex $< p_{start}(e),\ pos_{Sub} >$ to vertices $< p_{end}(e),\ pos_{Sub} + c >$ for all values of $c$ in a single call to the Edlib library [1]. Due to restrictions of Edlib library this implementation only supports edit distance ($\mu$ and $\sigma$ are equal to 1). SPAligner further enhance the optimal alignment finding approach presented above by various heuristics listed in Supplement Section "Alignment extension heuristics and thresholds".

#### 5 Generation of simulated reads and assembly graphs

Reference and dataset availability information is summarized in Table S1.
Assembly graphs were generated by SPAdes-3.12.0 [6] assembler with parameter -k 21,33,55,77.

Synthetic PacBio reads were generated by Pbsim [7] with option −−model_qc data/model_qc_clr. Synthetic Nanopore reads were generated by NanoSim [8] with default parameters and option circular for *E. coli* and linear for *C. elegans* and *S. cerevisiae.* Synthetic Illumina reads for *C. elegans* were generated by ART-MountRainier-2016-06-05 [9].

| Species | Reference | Illumina reads | PacBio reads | Nanopore reads |
|---|---|---|---|---|
| *E. coli* K12 | U00096.2 | ERA000206 | PacificBiosciences DevNet | Loman Lab R9 data |
| *S. cerevisiae* S288C | GCA_000146045.2 | ERP016443 | ERR1655118 | ERP016443 |
| *C. elegans* Bristol N2 | GCA_000002985.3 | – | PacificBiosciences DevNet | PRJEB22098 |

**Supplementary Table S 1**: Due to lack of freely available up-to-date ONT data for strain Bristol N2, reads sequenced from closely-related wild-type *C. elegans* strain VC2010 [10] were used (see original study for discussion of their relatedness).

#### 6 Notes on running aligners

Full archive with benchmarking data and scripts can be uploaded from https://figshare.com/s/b0dc3f715e0224e3e962
**SPAligner.** SPAligner accepts query sequences in fastq/fasta format and assembly graph in the GFA format. While edge-labeled assembly graphs considered in this paper are not expressive enough to represent an arbitrary graph in GFA format, assembly graphs produced by popular de Bruijn graph based assemblers (e.g. SPAdes and Megahit) can be considered as edge-labeled without loss of information.

Resulting alignments are output in custom tsv and/or GPA and fasta formats. In our benchmarks SPAligner was run with default settings.

**vg.** Assembly graphs were first converted to vg format by the vg mod -X 1024 command. For long read alignment we used the appropriate mode of the vg map command (vg map -m long), leaving other parameters to defaults. For each query sequence vg provides a list of non-overlapping local alignments ordered by their position on a read. Resulting local alignments were further processed to infer the longest continuous path $P$ consisting of edges $e_1, \ldots, e_n$

with consecutive local alignments ($P$ starts from the beginning of the first local alignment on $e_1$ and ends on the final position of the last local alignment on $e_n$).

**GraphAligner.** GraphAligner accepts assembly graphs in GFA format and outputs alignment in custom json format. GraphAligner was run with default settings. For *E. coli* datasets we additionally ran it with try-all-seeds flag recommended by the developers for bacterial data.

#### 7 Supplementary Table S2

| Aligner | MR(%) | AI (%) | T (days h:m) | M (Gb) | MR(%) | AI (%) | T (days h:m) | M (Gb) |
|---|---|---|---|---|---|---|---|---|
| *E. coli* PacBio | | | | | *E. coli* ONT | | | |
| vg | 9 | 83 | 00:16 | 3.9 | 68 | 87 | 01:12 | 4.2 |
| GraphAligner | 99 | 83 | 00:01 | 0.1 | 96 | 88 | 00:01 | 0.1 |
| GA try-all-seeds | 99 | 83 | 00:01 | 0.1 | 96 | 88 | 01:57 | 7 |
| SPAligner | 99 | 83 | 00:01 | 0.2 | 96 | 88 | 00:02 | 0.4 |
| *S. cerevisiae* PacBio | | | | | *S. cerevisiae* ONT | | | |
| vg | 8 | 83 | 07:55 | 90 | 21 | 83 | 14:13 | 92 |
| GraphAligner | 98 | 83 | 00:01 | 0.2 | 84 | 83 | 00:01 | 0.2 |
| SPAligner | 98 | 83 | 00:10 | 0.6 | 85 | 83 | 00:20 | 0.7 |
| *C. elegans* PacBio | | | | | *C. elegans* ONT | | | |
| vg | 7 | 83 | 1 day 03:51 | 314 | 20 | 88 | 3 days 15:26 | 301 |
| GraphAligner | 99 | 84 | 00:01 | 1.3 | 80 | 88 | 00:03 | 1.7 |
| SPAligner | 98 | 83 | 0:16 | 1.1 | 88 | 88 | 00:33 | 1.5 |

**Supplementary Table S 2**: Summary statistics of aligning synthetic PacBio/ONT reads to short-read assembly graphs (constructed by SPAdes with k-mer size equal to 77). Each dataset consists of 10k reads longer than 2 Kbp. All runs performed in 16 threads. While the number of mapped reads increases by 10-15% (as compared to real datasets) across all tools, similar conclusions about tools efficiency can be made. Specifically, SPAligner showed the best performance with respect to the number of mapped reads (for ONT datasets) and memory, GraphAligner showed the best performance with respect to speed, and SPAligner and GraphAligner showed roughly the same performance with respect to average identity.

#### 8 Supplementary Table S3

| | *E. coli* PacBio(%) | *E. coli* ONT (%) | *E. coli* sim PacBio (%) | *E. coli* sim ONT (%) |
|---|---|---|---|---|
| No. of mapped reads | 84 | 80 | 99 | 96 |
| Avg identity GraphAligner | 88 | 87 | 83 | 88 |
| Avg identity SPAligner | 88 | 87 | 83 | 88 |
| | *S. cerevisiae* PacBio(%) | *S. cerevisiae* ONT (%) | *S. cerevisiae* sim PacBio (%) | *S. cerevisiae* sim ONT (%) |
| No. of mapped reads | 57 | 60 | 98 | 84 |
| Avg identity GraphAligner | 88 | 83 | 83 | 83 |
| Avg identity SPAligner | 87 | 83 | 83 | 83 |

**Supplementary Table S 3**: Summary statistics on reads mapped by both SPAligner and GraphAligner. The table was generated by taking the intersection of sets of reads that were successfully mapped by SPAligner and GraphAligner. For each dataset and tool we identified percent of such reads and average identity of the resulting alignments.

#### References


[1] Sosic M, Sikic M. Edlib: a C/C++ library for fast, exact sequence alignment using edit distance;33(9):1394–1395. Available from: https://www.ncbi.nlm.nih.gov/pubmed/28453688.

[2] Amir A, Lewenstein M, Lewenstein N. Pattern Matching in Hypertext;35(1):82–99. Available from: https://linkinghub.elsevier.com/retrieve/pii/S0196677499910635.

[3] Antipov D, Korobeynikov A, McLean JS, Pevzner PA. hybridSPAdes: an algorithm for hybrid assembly of short and long reads;32(7):1009–1015. Available from: http://dx.doi.org/10.1093/bioinformatics/btv688.

[4] Navarro G. A guided tour to approximate string matching;33(1):31–88. Available from: http://portal.acm.org/citation.cfm?doid=375360.375365.

[5] Rautiainen M, Marschall T. Aligning sequences to general graphs in ( + ) time;Available from: http://biorxiv.org/lookup/doi/10.1101/216127.

[6] Nurk S, Bankevich A, Antipov D, Gurevich A, Korobeynikov A, Lapidus A, et al. Assembling Genomes and Mini-metagenomes from Highly Chimeric Reads. In: Deng M, Jiang R, Sun F, Zhang X, editors. Research in Computational Molecular Biology. vol. 7821. Springer Berlin Heidelberg;. p. 158–170. Available from: http://link.springer.com/10.1007/978-3-642-37195-0_13.

[7] Ono Y, Asai K, Hamada M. PBSIM: PacBio reads simulatortoward accurate genome assembly;29(1):119–121. Available from: http://dx.doi.org/10.1093/bioinformatics/bts649.

[8] Yang C, Chu J, Warren RL, Birol I. NanoSim: nanopore sequence read simulator based on statistical characterization;6(4):gix010–gix010. Available from: http://dx.doi.org/10.1093/gigascience/gix010.

[9] Huang W, Li L, Myers JR, Marth GT. ART: a next-generation sequencing read simulator;28(4):593–594. Available from: http://dx.doi.org/10.1093/bioinformatics/btr708.

[10] Tyson JR, O'Neil NJ, Jain M, Olsen HE, Hieter P, Snutch TP. MinION-based long-read sequencing and assembly extends the Caenorhabditis elegans reference genome;28(2):266–274. Available from: https://www.ncbi.nlm.nih.gov/pubmed/29273626.
